# Organic stabilization of extracellular elemental sulfur in a *Sulfurovum*-rich biofilm: a new role for EPS?

**DOI:** 10.1101/2021.06.03.446708

**Authors:** Brandi Cron, Jennifer L. Macalady, Julie Cosmidis

## Abstract

This work shines light on the role of extracellular polymeric substances (EPS) in the formation and preservation of elemental sulfur biominerals produced by sulfur-oxidizing bacteria. We characterized elemental sulfur particles produced within a *Sulfurovum*-rich biofilm in the Frasassi Cave System (Italy). The particles adopt spherical and bipyramidal morphologies, and display both stable (α-S_8_) and metastable (β-S_8_) crystal structures. Elemental sulfur is embedded within a dense matrix of EPS and the particles are surrounded by organic envelopes rich in amide and carboxylic groups. Organic encapsulation and the presence of metastable crystal structures are consistent with elemental sulfur organomineralization, i.e. the formation and stabilization of elemental sulfur in the presence of organics, a mechanism that has previously been observed in laboratory studies. This research provides new evidence for the important role of microbial EPS in mineral formation in the environment. We hypothesize that extracellular organics are used by sulfur-oxidizing bacteria for the stabilization of elemental sulfur minerals outside of the cell wall as a store of chemical energy. The stabilization of energy sources (in the form of a solid electron acceptor) in biofilms is a potential new role for microbial EPS that requires further investigation.

## 1 Introduction

Elemental sulfur (S(0)) is an intermediate of the biogeochemical sulfur cycle found in many natural environments such as marine sediments, marine and lacustrine water columns, cold or hot springs, hydrothermal environments, salt marshes and caves (Findlay et al., 2014; Hamilton et al., 2015; Jørgensen et al., 2019; Lau et al., 2017; Zerkle et al., 2010). S(0) is formed by chemical or biological oxidation of more reduced sulfur species, although in low-temperature environments biological S-oxidation rates are typically more than three orders of magnitude faster than abiotic rates (Luther et al., 2011). A wide diversity of microorganisms can oxidize sulfide, polysulfides, or thiosulfate and precipitate S(0) through both phototrophic and chemotrophic pathways (Dahl and Prange, 2006; Kleinjan W.E., 2003). In turn, microbially formed S(0) can be used as a source of energy for a wide diversity of S-oxidizers, S-reducers, and microorganisms that perform S(0) disproportionation (Dahl, 2020a; Warthmann et al., 1992). Elemental sulfur thus occupies a central and ecologically important role in the biogeochemical sulfur cycle.

Biogenic S(0) is deposited either intracellularly or extracellularly (Dahl and Prange, 2006; Kleinjan W.E., 2003; Maki, 2013). Most studies on microbial S(0) biomineralization so far have focused on microorganisms forming intracellular S(0) globules, for instance *Allochromatium vinosum, Acidithiobacillus ferrooxidans, Thiothrix* spp., or large colorless SOB such as *Thiomargarita namibiensis, Thioploca* spp. or *Beggiatoa* spp. (Brune, 1995; Dahl, 2020b; Gray and Head, 1999; Maki, 2013; Nims et al., 2019; Pasteris et al., 2001; Prange et al., 2002). Many SOB can form S(0) extracellularly, such as green sulfur bacteria (Chlorobiaceae) (Gregersen et al., 2011; Marnocha et al., 2016), purple sulfur bacteria of the Ectothiorhodospiraceae family (Then and Trüper, 1983), some purple non-sulfur bacteria (Hansen and van Gemerden, 1972), some lithotrophic sulfur bacteria (Cron et al., 2019) and cyanobacteria (Oren and Shilo, 1979). More work is needed to decipher the formation mechanisms of extracellular S(0) and to characterize its properties.

Elemental sulfur S(0) is thermodynamically unstable under a wide range of natural Eh-pH conditions (especially at circumneutral pH; Rickard and Luther, 2007), so it is not clear how microbial sulfur, particularly when it is extracellular, persists in the environment. Cosmidis et al. (2019) recently showed that interactions with organics are important for the abiotic formation of S(0) minerals, a process termed S(0) organomineralization. Organomineralized sulfur can be found in several metastable crystal structures including the monoclinic allotropes β-S_8_ and γ-S_8_, which are thought to be stabilized by close association with organics. Organic-mineral interactions may also be important in extracellular S(0) formation by bacteria. Indeed, organics produced by the chemolithoautotrophic SOB *Sulfuricurvun kujiense* (Campylobacterota) are needed for extracellular S(0) formation by this organism. The S(0) globules of *S. kujiense* are composed of β-S_8_ and γ-S_8_ and are coated by organic envelopes that allow them to precipitate under conditions outside of their theoretical thermodynamic stability domain (Cron et al., 2019). Other SOB such as *Thiobacillus* sp. W5 (Kleinjan et al., 2005) and *Chlorobaculum tepidum* (Hanson et al., 2016; Marnocha et al., 2019) also produce extracellular S(0) globules with organic envelopes, suggesting that organics play a previously overlooked role in microbial S(0) formation and stabilization in nature.

Previous studies describing the importance of organics in S(0) mineralization were based on laboratory experiments, whereas observations from natural environments are still lacking. Elemental sulfur particles with metastable structures and intimate associations with organics were described in a sulfur-rich glacial site in the Arctic, but it could not be determined whether microbial mediation was involved in their formation (Lau et al., 2017). In the present study, we characterized extracellular S(0) particles formed within microbial biofilms in a subsurface environment dominated by sulfur-cycling bacteria. The Grotto Grande del Vento-Grotta del Fiume (Frassasi) cave system is actively forming in Jurassic limestone in the Apennine Mountains of the Marches Region, Central Italy (D’Angeli et al., 2019). The S(0) minerals described here are found within microbial biofilms in a microaerophilic sulfide-rich stream, Pozzo di Cristalli. Previous full-cycle rRNA and metagenomic approaches identified Campylobacterota most closely affiliated with *Sulfurovum* species as the most abundant organisms in streamer biofilms at this location (Hamilton et al., 2015; Jones et al., 2008; Macalady et al., 2008). *Sulfurovum* oxidize sulfide and thiosulfate to sulfate and extracellular S(0) globules, which can make up more than 60% by weight of their mats. These primary producers serve as the principal source of organic carbon to the subsurface stream ecosystem (Hamilton et al., 2015). The biofilms form attached to rocks in the stream bed where they intersect the water surface. In this environment, sulfidic water turbulently mixes with oxygen in the cave air, providing chemical energy for growth (Macalady et al., 2008).

We characterized the morphology and crystal structure of S(0) particles formed in these natural *Sulfurovum*-dominated biofilms using Scanning Electron Microscopy (SEM), X-Ray Diffraction (XRD) and Ultra-low frequency Raman spectromicroscopy. Organics closely associated with the S(0) minerals were also characterized using Fourier Transform Infrared (FTIR) spectroscopy and Scanning Transmission X-ray Microscopy (STXM). Our results provide important clues about how extracellular polymeric substances (EPS) stabilize extracellular S(0) in the environment. We discuss the ecological implications of this new role of EPS in sulfur-based microbial ecosystems.

## 2 Methods

### 2.1 Biofilm sample collection

Pozzo di Cristalli is a cave stream that periodically hosts blooms of streamer biofilms primarily composed of Campylobacterota in the genus *Sulfurovum* (Hamilton et al., 2015; Jones et al., 2008; Macalady et al., 2008). The biofilms thrive in turbulent stream riffles near the air-water interface. Sulfide concentration was 20.5 μM at the time of sampling in September 2016. Streamer sample PC1647 was harvested from customized floating twine supports (Henri et al., *in prep.*). In September 2017, a *Sulfurovum*-dominated biofilm (streamer sample PC1718) was collected from a limestone cobble at the same location (Fig. 1). Biofilm samples were divided into sterile Falcon tubes and cryotubes. Samples in Falcon tubes were preserved with glutaraldehyde and stored at −20 °C. Cryotube samples were stored at −80 °C.

**Figure 1:**
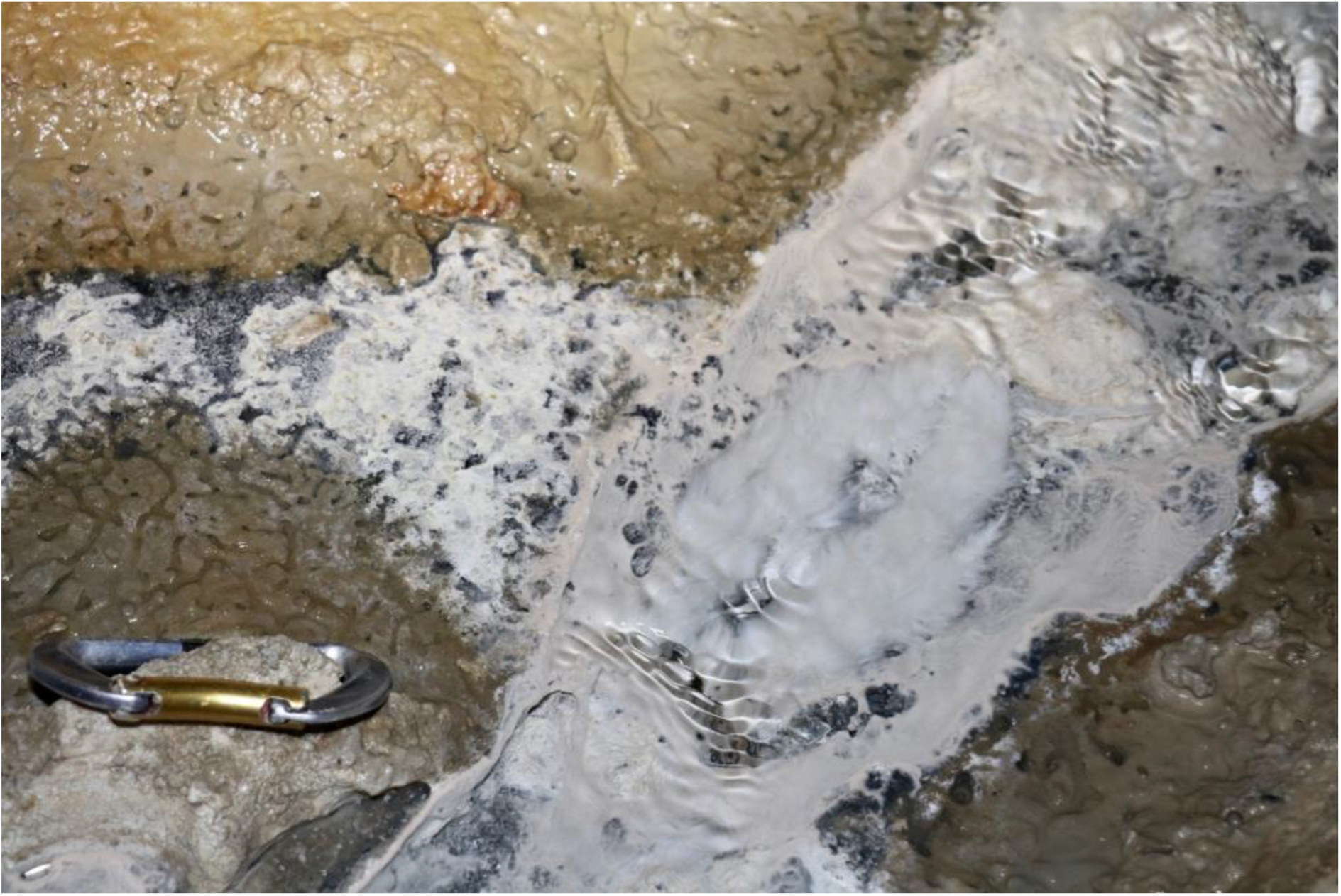
Biofilm collection site in September 2017 with billowing white *Sulfurovum*-rich streamer biofilm on the surface of the stream at center right. The *Sulfurovum*-dominated biofilm is surrounded by a weakly pigmented *Beggiatoa* biofilm growing on the black sediment surface.

### 2.2 X-ray diffraction

For X-ray Diffraction (XRD), 1 mL of biofilm (sample PC1718) was rinsed three times with deionized water and deposited on a single crystal miscut Si holder. Samples were analyzed using a PANalytical Empyrean diffractometer paired with a PIXcel3D detector, and Cu Kα (λ = 1.5406 Å) incident X-ray radiation. Scans were conducted over a 2θ range of 5–70°, and analyses used a step size of 0.025°, a time of 96.4 seconds per step, and a current density of 40 mA. XRD data were analyzed using MDI JADE software.

### 2.3 Raman spectromicroscopy

For Raman, 5 mL of frozen unpreserved biofilm (sample PC1718) were rinsed 3 times with deionized water to remove salts. Samples were either deposited and dried on a microscope slide or kept wet between a microscope slide and cover slip. Raman spectra were collected using a Horiba LabRam HR Evolution Vis-NIR optimized & AIST-NT Scanning Probe, and a Si-based CCD detector (1024 × 256 pixels). Raman signals were measured in the low frequency range using BragGrate notch filters (Nims et al., 2019). The spectrometer was calibrated using the 520 cm^−1^ Raman peak of Si prior to analysis. Spectral data were corrected for instrumental artifacts and baseline-subtracted using a polynomial fitting algorithm in LabSpec 6 (Horiba Scientific). The sample spectra were compared with reference Raman spectra for different allotropes of S(0) (Nims et al., 2019).

### 2.3 Scanning Electron Microscopy

For Scanning Electron Microscopy (SEM), biofilm samples were rinsed with deionized water and deposited on polycarbonate filters (GTTP Isopore membrane filters, Merck Millipore, pore size 0.2 μm) or on glass slides (for correlative Raman analyses, see below). The samples were allowed to dry at ambient temperature and coated with iridium or gold prior to analysis. For sample PC1718, SEM analyses were conducted on a Field Emission Nova NanoSEM 630 at the Materials Characterization Laboratory at The Pennsylvania State University. Images were acquired with the microscope operating at 3-7 kV and a working distance (WD) of ~3-5 mm. Energy-Dispersive X-ray Spectroscopy (EDXS) analyses were performed at 12 kV and WD ~7 mm. For sample PC1647, SEM analyses were conducted on a JSM-7401F field emission scanning electron microscope (FESEM) at the Nanoscale Fabrication Laboratory at the University of Colorado at Boulder. Images were acquired in the secondary electron mode with the microscope operating at 5 kV and a WD of 6 mm, and in the backscattered electron mode at 15 kV and WD 8 mm. EDXS analyses were performed at 20 kV and WD 8 mm.

To correlate Raman data with morphological characterization of the S(0) particles, we designed a correlative Raman-SEM protocol. A frozen unpreserved biofilm sample (PC1718) was rinsed 3 times with deionized water to remove salts, and deposited on a microscope glass slide. The sample was air-dried, and the S(0) particles were first analyzed using Raman. The microscope slide was then coated with iridium and analyzed by Scanning Electron Microscopy (SEM) on a Field Emission Nova NanoSEM 630 (see details on SEM operations below). Raman-based crystallographic characterization of the S(0) minerals was performed before SEM imaging in order to prevent potential structural alteration in the dry, low-vacuum environment of the SEM chamber.

### 2.4 Fourier-Transform Infrared Spectroscopy

For Fourier-Transform Infrared Spectroscopy (FTIR), 1 mL of unpreserved biofilm (sample PC1718) was rinsed three times with deionized water and dried in a vacuum oven at 60 °C overnight. Then, 6 mg of dried sample and 100 mg KBr were ground and pelleted. FTIR measurements were conducted on a vertex 70 spectrometer (Bruker Optics) equipped with a deuterated triglycerine sulfate (DTGS) detector and a high intensity water cooled Globar source. Spectra were collected at 5 cm^−1^ resolution (2.5 mm aperture) as an average of 100 scans using MVP PRO software (Harrick Scientific). The instrument was purged for 30 minutes before the first measurement to ensure baseline stability. The spectra were baseline corrected using the ‘Rubber Band’ algorithm within the OPUS 2.2 software. The experimental spectra were plotted with reference spectra for calcite and quartz (RRUFF database; Lafuente et al., 2015).

### 2.5 Scanning Transmission X-ray Microscopy and C K-edge and S L-edge X-ray absorption spectroscopy

For Scanning Transmission X-ray Microscopy (STXM), 1 mL of unpreserved biofilm (samples PC1647 and PC1718) was rinsed three times with deionized water. A small drop of the suspension (~3 μL) was deposited on a Formvar-coated 200 mesh Cu TEM grid (Ted Pella) and allowed to air-dry at ambient temperature. STXM analyses were performed on beamline 10ID-1 (SM) of the Canadian Light Source (Saskatoon, Canada), and beamline 11.0.2. of the Advanced Light Source (ALS, Berkeley, CA). The X-ray beam was focused on the samples using a Fresnel zone plate objective and an order-sorting aperture yielding a focused X-ray beam spot of ~30 nm on the samples. After sample insertion in the STXM microscope, the chamber was evacuated to 100 mTorr and back-filled with He at ~1 atm pressure. Energy calibration was achieved using the well-resolved 3p Rydberg peak of gaseous CO2 at 294.96 eV. Images, maps and image stacks were acquired in the 260-340 eV (C K-edge) and 155-190 eV (S L-edge) energy ranges.

STXM data were processed using aXis2000 software. A linear background correction was applied to the spectra at the C K-edge and the S L-edge, in the 260-280 eV region and 155-160 eV region, respectively, to eliminate the contributions of lower energy absorption edges. Maps of organic C were obtained by subtracting images obtained at 280 eV (i.e. below the C K-edge) and converted into optical density (OD) from OD-converted images at 288.2 eV (1s→π* electronic transitions in amide groups). Maps of S were obtained by subtracting OD-converted images obtained at 160 eV (i.e. below the S L-edge) from OD-converted images at 163.5 eV (energy of the S L3-edge). X-ray absorption near edge structure (XANES) spectra were extracted from image stacks as explained in Cosmidis and Benzerara (2014).

## 3 Results

### 3.1 Morphology of elemental sulfur particles in *Sulfurovum*-dominated biofilms

SEM imaging combined with EDXS analyses of the *Sulfurovum*-dominated biofilms revealed abundant sulfur-rich particles within a dense matrix of extracellular polymeric substances (EPS) (Figs. 2,S1,S2). The particles appear as spheroids and bipyramidal crystals, which are sometimes fused into elongated chains (Figs. 2A, S1). The particles may also adopt more irregular shapes. The EPS appear either film-like (Fig. 2C) or as a web made from a network of thin threads (Fig. 2F). Figure 3 shows the size distribution of S(0) spheroids in two biofilm samples (PC1647 (n=1354) and PC1718 (n=256)). The diameters of the spheroids range from 0.1 to 3.5 μm, with an average diameter of 1.31 μm for PC1347 and 1.03 μm for PC1718. Bipyramids are not abundant enough to plot their size distribution, but range between 1.9 and 4.8 μm in length.

**Figure 2:**
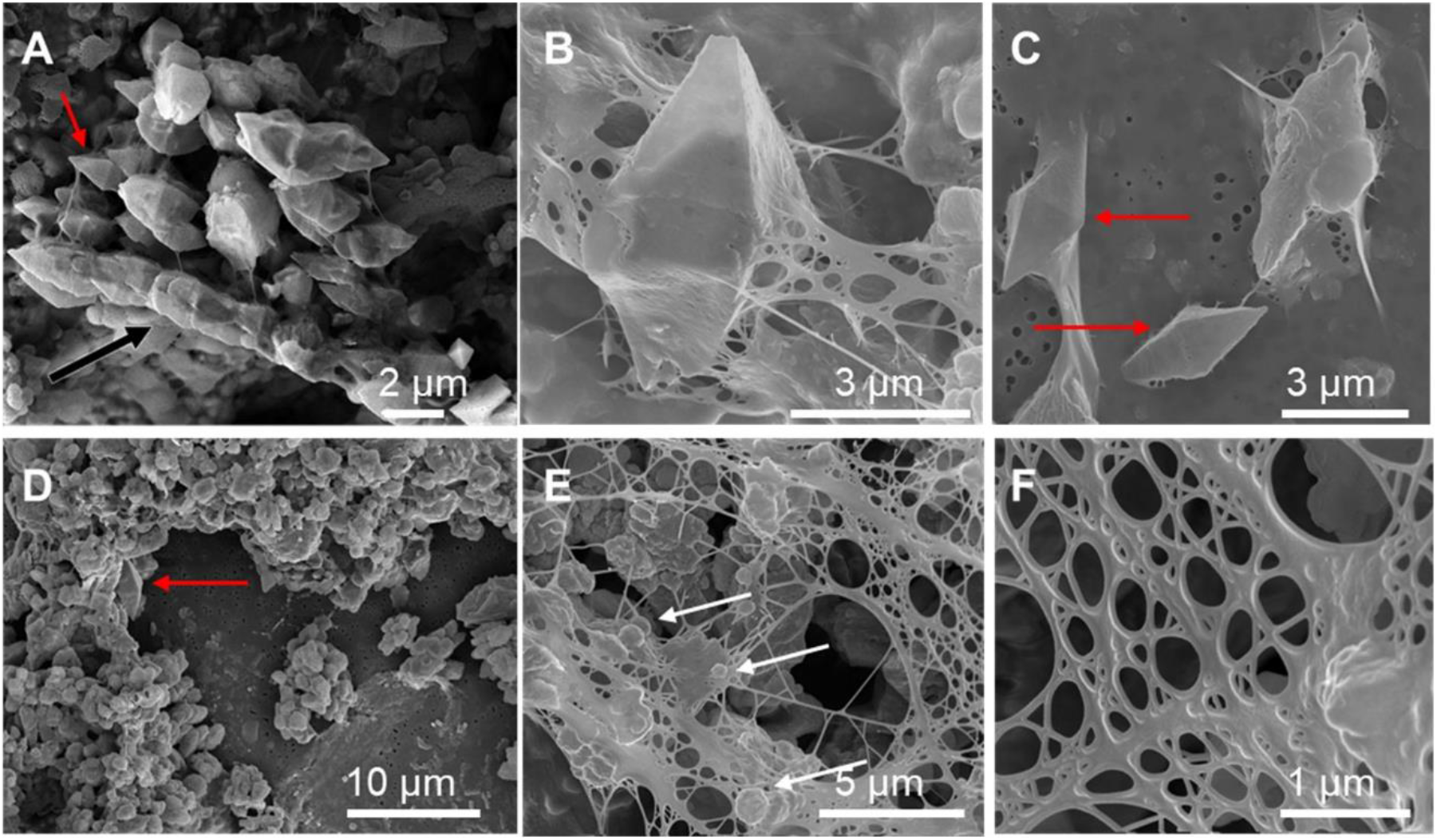
Scanning electron microscopy images of sulfur particles associated with EPS in *Sulfurovum*-dominated streamer biofilms. Samples include (A) PC1647 and (B-F) PC1718. Red arrows point to bipyramidal S(0) crystals and white arrows point to S(0) spheroids. (A) shows an example of a chain formed by fused bipyramidal S(0) crystals (black arrow).

**Figure 3:**
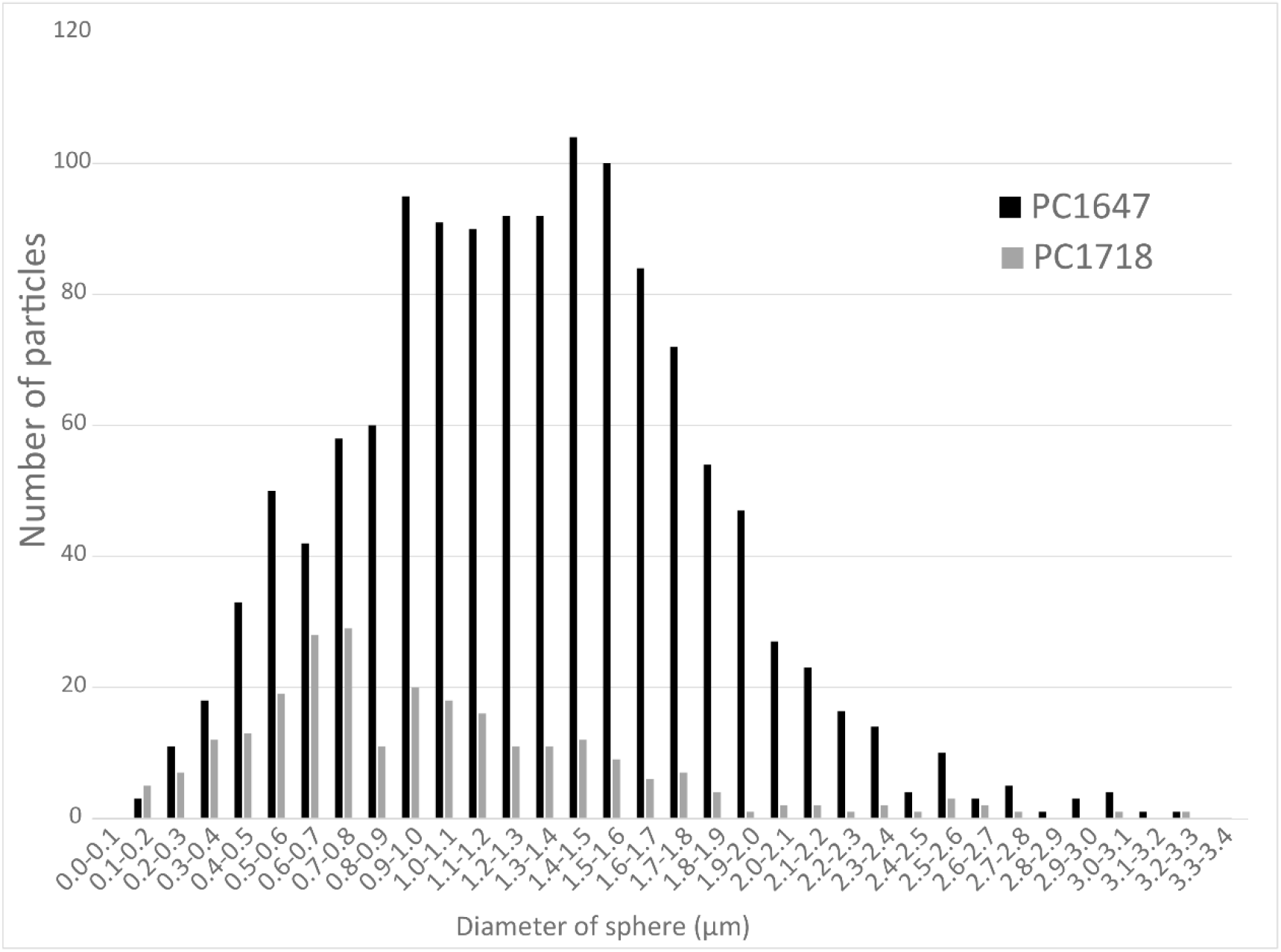
Histograms depicting the size distributions of spherical S(0) particles in the PC1647 and PC1718 biofilm samples.

### 3.2 Crystal structure of elemental sulfur

#### 3.2.1 X-ray diffraction

XRD analyses performed on sample PC1718 indicate that the biofilm contains S(0) present as orthorhombic α-S_8_ and monoclinic β-S_8_ (Fig. 4). Calcite and quartz minerals were also detected.

**Figure 4:**
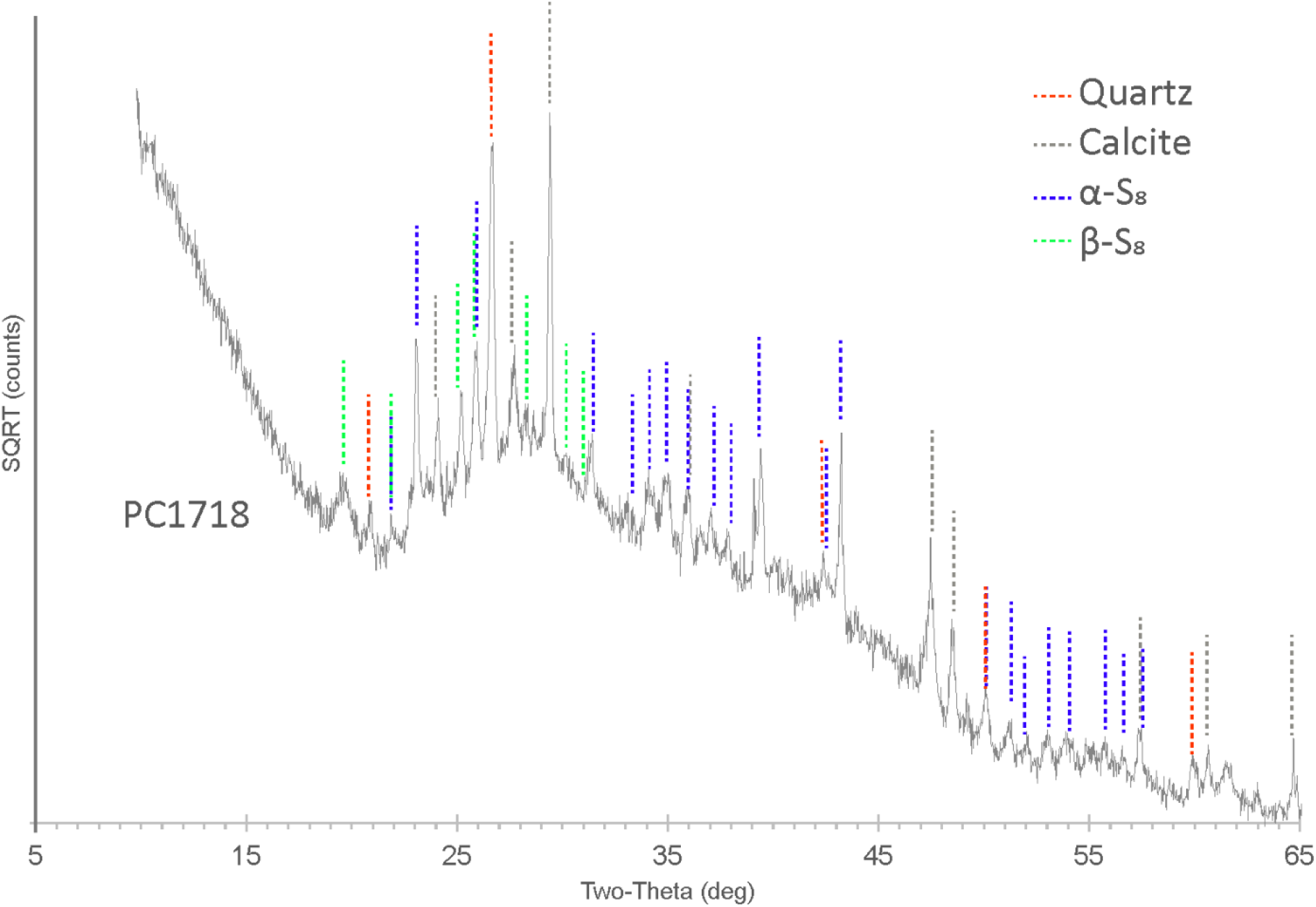
XRD spectrum for biofilm sample PC1718. The peaks in the diffractogram include matches for α-S_8_, β-S_8_, quartz, and calcite.

#### 3.2.2 Correlative scanning electron microscopy and Raman spectromicroscopy

In order to determine the relationship between S(0) crystal structure and particle morphology, Raman spectroscopy correlated with SEM imaging was performed on sample PC1718. The α-S_8_ allotrope can be identified using the low frequency range Raman vibration modes at 28 cm^−1^, 44 cm^−1^, 51 cm^−1^, 63 cm^−1^, and a doublet at 82 cm^−1^ and 88 cm^−1^ (Fig. 5C). The β-S_8_ allotrope displays distinctive vibrations in the low frequency range with a peak positioned at 82 cm^−1^, a doublet at 33 cm^−1^ and 42 cm^−1^ and a shoulder at 60 cm^−1^. Spheroids were found to be composed of either α-S_8_ or β-S_8_ (Fig. 5A, B, D). These two allotropes were sometimes found in close association with each other (Fig. 5E). We attempted to determine the crystal structure of S(0) bipyramids but they could not be found in the sample during Raman analyses due to their relative rarity.

**Figure 5:**
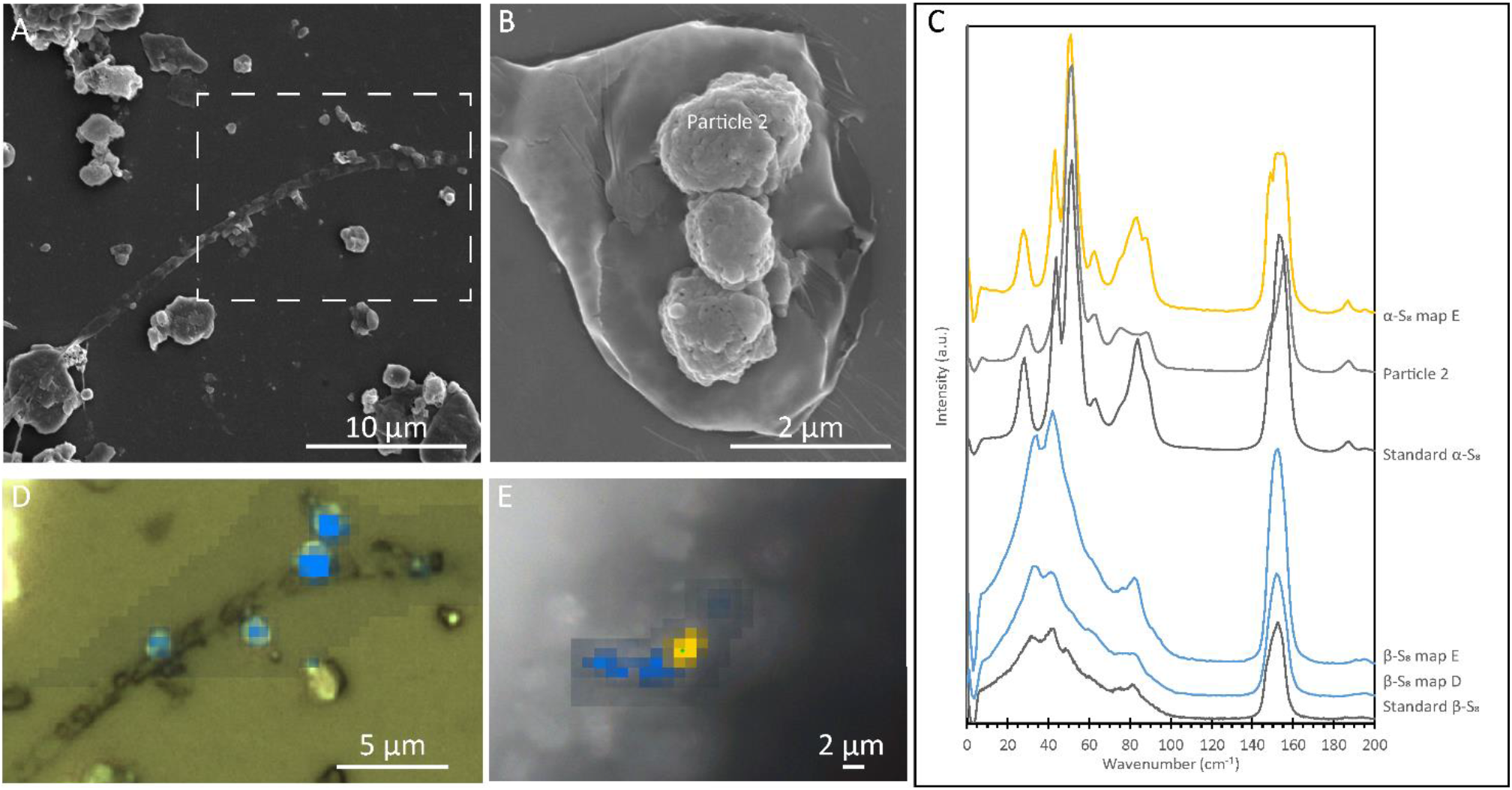
Correlative SEM and Raman analyses of a filamentous cell with extracellular S(0) in the *Sulfurovum*-dominated streamer biofilm (sample PC1718). The sample was airdried in images (A), (B), and (D), while Raman analyses (E) were performed while the sample was wet. (A) SEM image with a white dashed box indicating the location of the Raman map in (D); (B) SEM image of particles in sample PC1718. “α-S_8_ particle 2” indicates where the α-S_8_ Raman spectrum (labelled “Particle 2”) in (C) was collected. (C) Ultra-low frequency Raman spectra. Standard spectra for α-S_8_ and β-S_8_ are labelled accordingly. (D) Light microscopy image with an overlay Raman map. The blue areas are rich in β-S_8_. (E) light microscopy image with a Raman map overlayed. Yellow areas are rich in α-S_8_ and blue areas are rich in β-S_8_.

### 3.3 Association of elemental sulfur with EPS and encapsulating organics

#### 3.3.1 Fourier-Transform Infrared Spectroscopy

Fourier-Transform Infrared Spectroscopy (FTIR) was performed on the unpreserved PC1718 sample (Fig. 6). Reference spectra for calcite and quartz were used to assist in the interpretation of the FTIR data, since these minerals were identified in the sample based on XRD (Fig. 4). The sample spectrum displays a broad band centered around 3400-3430 cm^−1^ corresponding to O-H stretching frequencies (Fig. S3). The broad peak around 1750 cm^−1^ corresponds to stretching of COOH or COOR in carboxylic acids and aromatic esters (Artz et al., 2008). Signal in this region is also attributed to C=O stretching in esters and fatty acids (Schmitt and Flemming, 1998). Peak signal around 1650-1600 cm^−1^ is attributed to stretching of aromatic C=C or asymmetric C-O stretching in COO^−^ (carboxylates) (Artz et al., 2008; Tinti et al., 2015). The presence of carboxylic acids in our sample is further supported by the presence of a peak at 1426 cm^−1^ originating from symmetric C=O stretching and OH deformation in COOH from carboxylates or carboxylic acid structures (Thomas, 1972). The band at 1056 cm^−1^ is attributed to phosphates in nucleic acids (Orhan Yanıkan et al., 2020), and/or to quartz. A sharp peak at 879 cm^−1^ is present, commonly interpreted as out of phase ring stretching (ring ‘breathing’) of aromatics (Artz et al., 2008). Peaks at 470 and 424 cm^−1^ are attributed to S-S stretching in S_8_ (Meyer, 1976; Steudel and Eckert, 2003). The presence of quartz is confirmed by peaks at 692, 776, and 795 cm^−1^ and a shoulder at 1160 cm^−1^. The peak at 516 cm^−1^ is representative of calcite. Quartz peaks may overlap with the molecular vibrations of polysaccharides in the 1200-900 cm^−1^ region, which are typically observed in FTIR spectra of EPS (Naumann et al., 1991).

**Figure 6:**
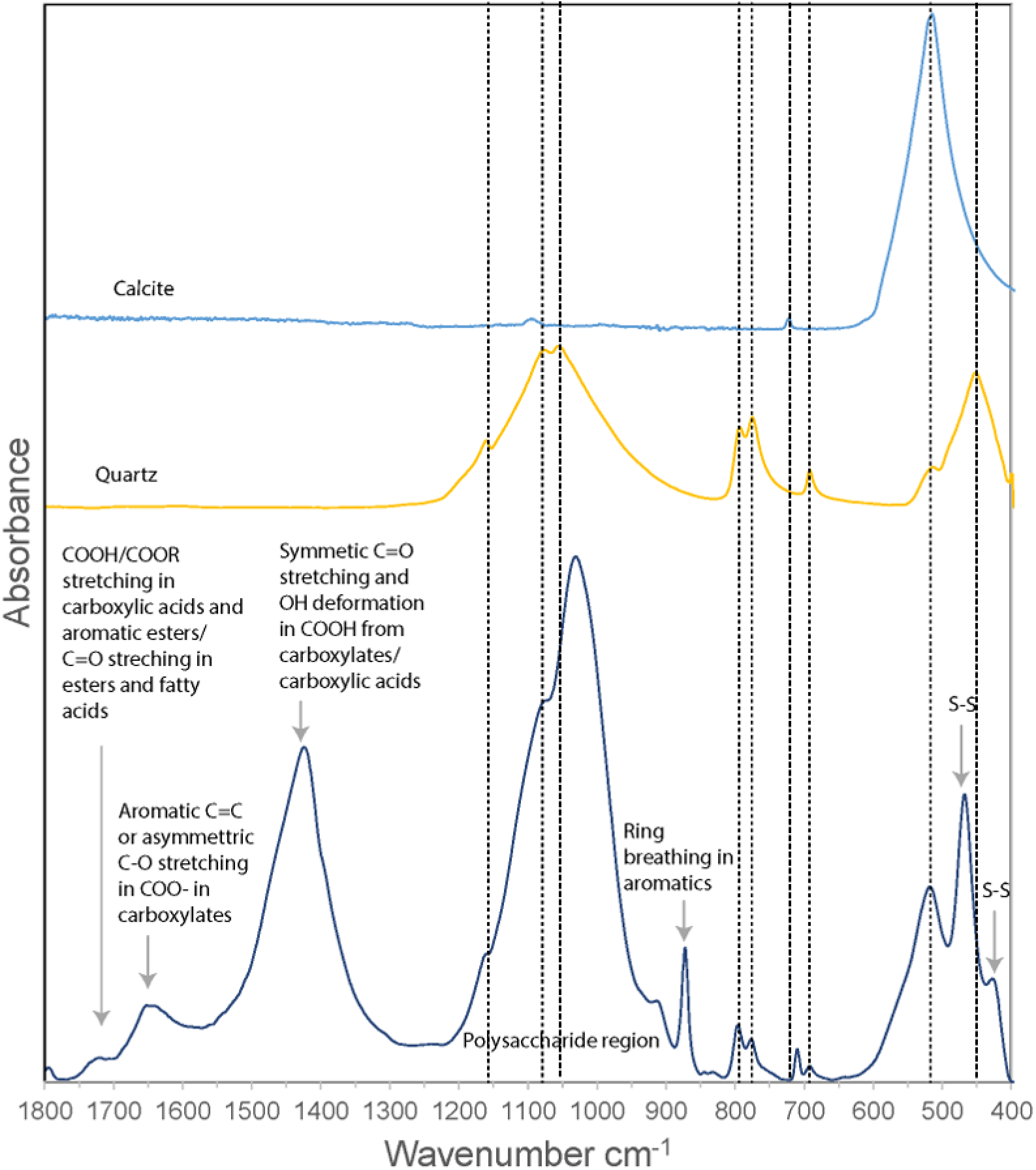
FTIR spectrum of *Sulfurovum*-dominated streamer biofilm (sample PC1718), plotted with reference spectra for calcite and quartz. Dashed vertical lines indicate the positions of the peaks corresponding to calcite and quartz.

FTIR analysis thus confirmed the presence of quartz, calcite, and S(0) minerals in the biofilm, while signal from the organic material shows a composition dominated by carboxylic acids, carboxylates, and aromatic structures.

#### 3.3.2 Scanning Transmission X-ray Microscopy at the C K-edge and S L-edge

STXM analyses at the S L-edge of the *Sulfurovum-*dominated biofilms (samples PC1647 and PC1718) confirmed that S(0) is the only form of particulate sulfur present in the samples (Fig. S4). C K-edge analyses showed the presence of bacteria along with two types of extracellular carbon materials: abundant diffuse EPS, and thin envelopes around S(0) particles (Fig. 7). The thin organic envelopes are especially visible in STXM images and maps in places where S(0) particles were vaporized due to the low pressure of the STXM chamber and X-ray beam damage (black arrows in Fig. 7A).

**Figure 7:**
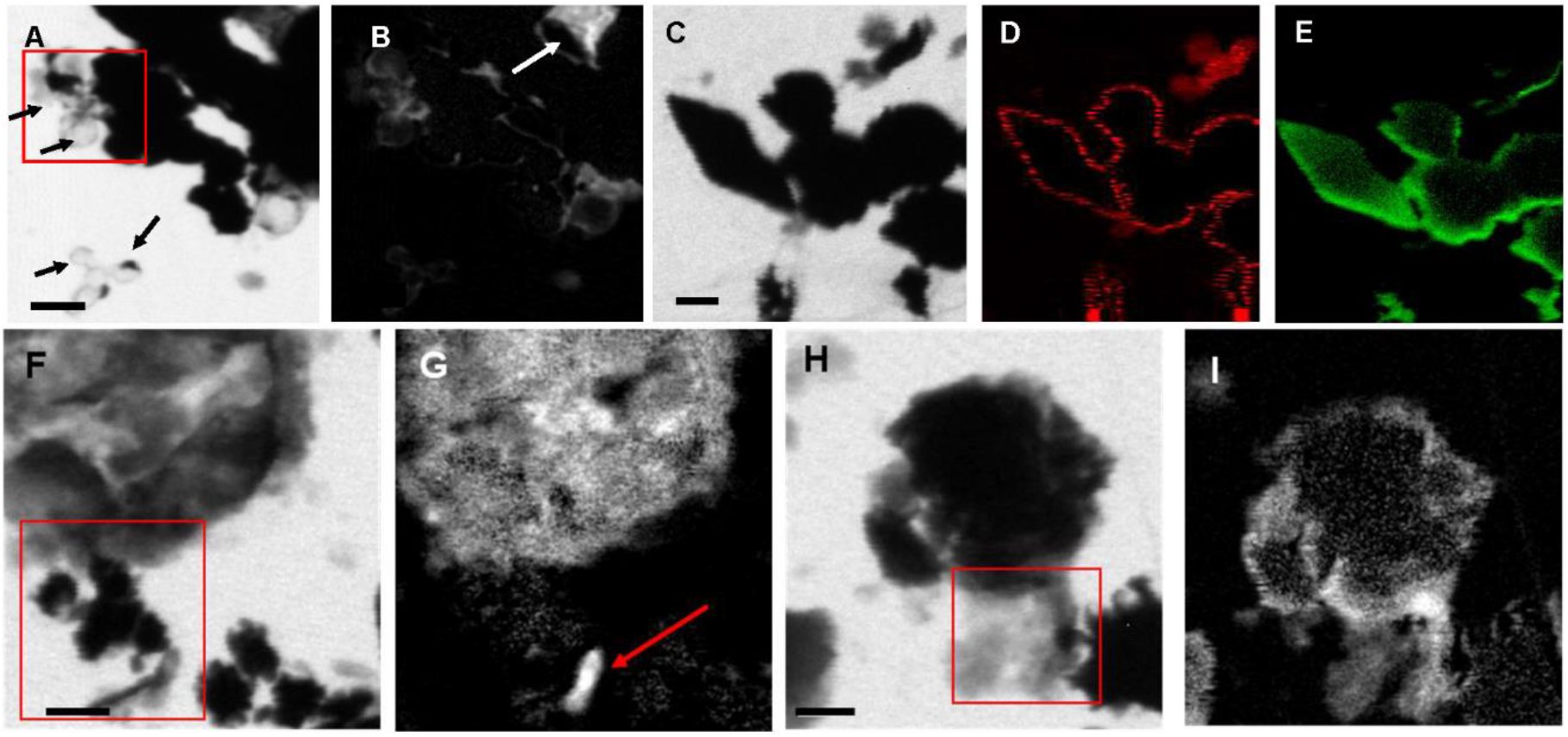
STXM images and maps of S*ulfurovum*-dominated streamer biofilm. (A-B) Sample PC1647. (A) image collected at 288.1 eV. Black areas are S(0) particles. The empty spherical envelopes of vaporized S(0) globules are visible (black arrows). (B) Corresponding carbon map. The white arrow shows the carbon enveloped around a partially vaporized S(0) particle (a chain of S(0) bipyramids). (C-I) Sample PC1718. (C) Image collected at 288.2 eV. (D) Carbon map. (E) Sulfur map. (F) Image collected at 300 eV. (G) Carbon map. The arrow points to a rod-shaped microbial cell. (H-I) Image collected at 300eV. (I) Carbon map. The boxes in (A) (F) and (H) and the arrow in (B) indicate where XANES analyses shown in Figure 8 were performed. Scale bars: 1 μm.

All C K-edge XANES spectra display peaks at 285-285.2 eV, characteristic of 1s→π* C=C transitions in either aromatic or unsaturated carbon (Haberstroh et al., 2006; Lehmann, 2009; Solomon, 2009) (Fig. 8). The main peak in all spectra is located at 288.2 eV, corresponding to 1s→π* transitions in amide groups of proteins (Benzerara et al., 2004; Chan et al., 2011; Haberstroh et al., 2006). The presence of amides is in contradiction with FTIR results which did not clearly detect this functional group. This discrepancy may be due to the fact that we focused our STXM analyses on clearly identifiable features such as microbial cells and organic envelopes around S(0) particles, while the signal from these small features may have been diluted by the signal of the more abundant EPS in bulk FTIR analyses. All spectra furthermore display a shoulder at 287.5 eV, corresponding to 3s→σ* transitions in aliphatics (Haberstroh et al., 2006; Lehmann, 2009), and a small peak at 289.4 eV, corresponding to 1s→3p/σ* transitions in hydroxylic groups (Brandes et al., 2004). Some spectra (mostly EPS) have a peak at 288.5 eV, attributed to the 1s→π*C=O electronic transitions in carboxylics (Boyce et al., 2002; Chan et al., 2011; Cody et al., 1998). A small peak at 286.6 eV, representative of 1s→π* transitions in phenolic groups, ketones, carboxylates, or aldehydes (Cosmidis et al., 2019; Lehmann, 2009; Moffet et al., 2011; Myneni, 2002) is only present in the organic envelope of a S(0) chain (Figs. 7B,8). Spectra of EPS and S(0) envelopes sometimes display a shoulder at 288.7eV, interpreted as C 1s→π* C=O electronic signature of acidic polysaccharides (Chan et al., 2011; Toner et al., 2009).

**Figure 8:**
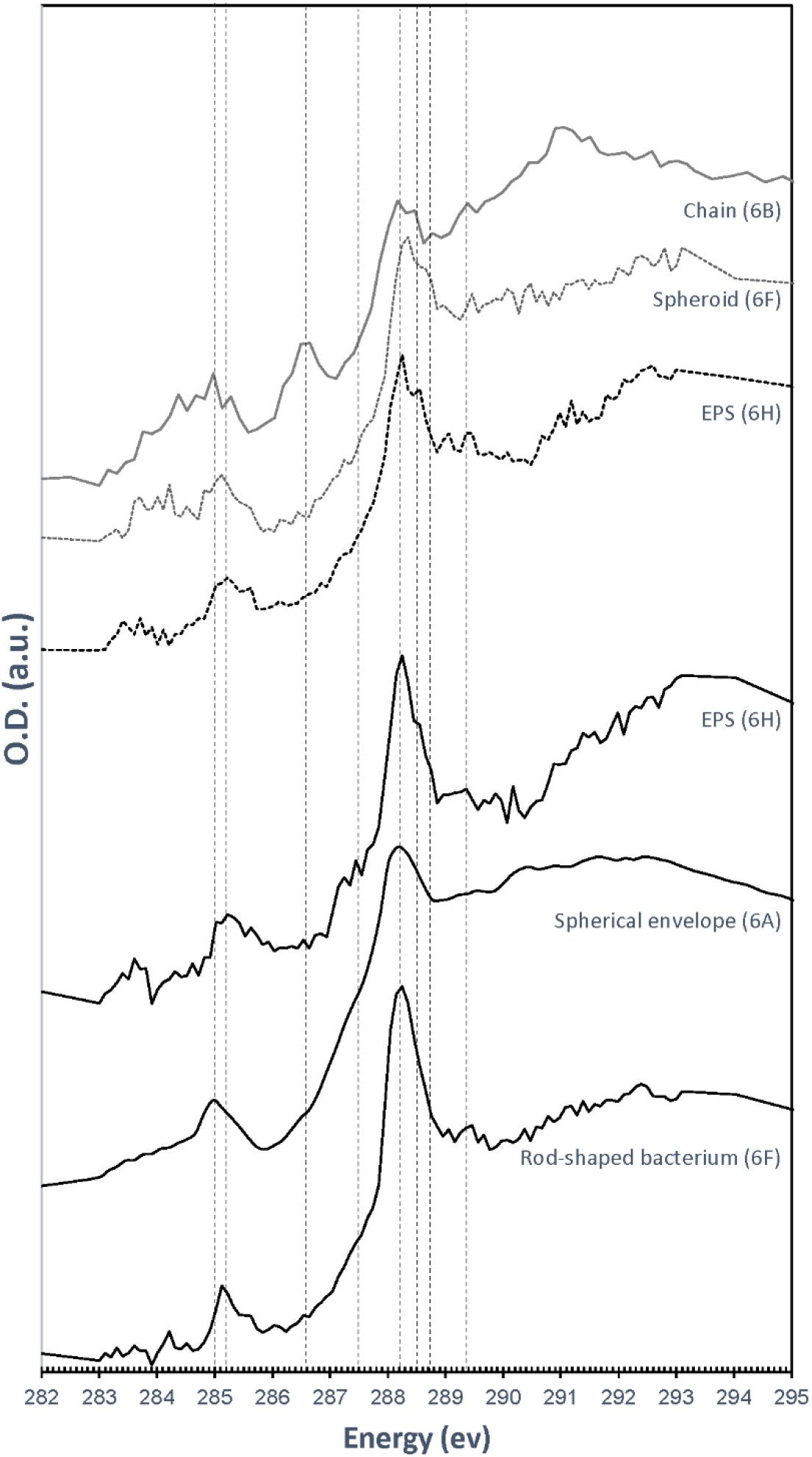
C K-edge XANES spectra obtained on the particles and cells shown in Figure 7. Dashed lines correspond to absorption energies for different organic functional groups: 285 eV (1s→π*C=C transitions in aromatics or unsaturated carbon), 285.2 eV (1s→π* C=C transitions in aromatics), 286.6 eV (1s→π* transitions in ketones, pyridines and phenols), 287.5 eV (3s→σ* transitions in aliphatics), 288.2 eV (1s→π* transitions in amides), 288.5 eV (1s→π*C=O transitions in carboxylic acids), 288.7 eV (1s→π* C=O transitions in carboxyls in acidic polysaccharides), and 289.4 eV (1s→3p/σ* transitions in alcohols, ethers and hydroxylated aliphatic compounds).

## 4 Discussion

### 4.1 Properties of the biofilm S(0) particles

We described S(0) particles forming within biofilms present in microaerophilic, sulfide-rich subsurface stream. The biofilms are dominated by Campylobacterota most closely realated to *Sulfurovum* species (Hamilton et al., 2015; Jones et al., 2008; Macalady et al., 2008), which are known to oxidize sulfide and thiosulfate to sulfate and/or extracellular S(0) (e.g., Campbell et al., 2006). Microbial sulfide oxidation rates are typically several orders of magnitude higher than chemical oxidation by molecular oxygen (Luther et al., 2011) suggesting that S(0) particles in the Frasassi biofilms are primarily the product of microbial S-oxidation. However, mineral nucleation and growth within biofilms is often influenced by surrounding EPS, which may result in particle shapes, sizes or crystal structures that differ from those of inorganically precipitated minerals (Tourney and Ngwenya, 2014). Here we summarize our observations of the S(0) particles in the Frasassi biofilms and suggest how formation within EPS may have influenced S(0) properties.

The samples we examined contained both S(0) spheroids and bipyramids, alongside more irregularly shaped particles (Figs. 2,7). Both spherical and bipyramidal morphologies have been observed in microbial biomineralization experiments, microbe-free organomineralization experiments, and inorganically precipitated S(0). Extracellular S(0) spheroids or globules are formed by diverse bacteria (Cron et al., 2019; Dahl and Prange, 2006; Marnocha et al., 2016). S(0) spheroids can also be formed through chemical precipitation of S(0) in the presence (Cosmidis et al., 2019) or in the absence (Marnocha et al., 2019) of organic compounds. Previous studies have suggested that bipyramids are typical of S(0) precipitated in the absence of organics (Steudel, 2003). However, as noted above, bipyramids have also been observed in S(0) organomineralization experiments with humic acids (Cosmidis et al., 2019) and in cultures of *C. tepdium* (Marnocha et al., 2019). The biofilm samples we examined contained S(0) spheroids ranging from 0.2 to 3.3 μm (Fig. 3) with median particle sizes near 1 μm. The PC1647 sample had larger S(0) spheroids. Larger particle size could be controlled by the age of the biofilm, or possibly by the growth of the biofilm on the twine. Further analysis would be needed to confirm whether biofilm age and the nature of attachment surfaces influence particle size. S(0) particles with sizes ranging from <0.2 μm to 10 μm have been observed in other natural environments (Findlay et al., 2014; Lau et al., 2017), in microbial cultures (Marnocha et al., 2019; Cron et al., 2019), in inorganically precipitated S(0) (Garcia and Druschel, 2014; Meyer, 1976), and in S(0) organomineralization experiments (Cosmidis et al., 2019). Particle sizes or morphologies of the S(0) described here are thus not particularly characteristic of their formation within microbial biofilms.

On the other hand, we observed both α-S_8_ and β-S_8_ S(0) crystal structures in biofilm sample PC1718 (Figs. 4,5). Sample PC1647, analyzed by Henri et al. (*in prep.*), contained only α-S_8_, which again could be due to biofilm aging or growth on a twine. The monoclinic sulfur allotrope β-S_8_ is thermodynamically unstable at temperatures lower than 96 °C, as opposed to orthorhombic α-S_8_ which is the stable structure at room temperature (Steudel, 2003). In laboratory studies, metastable monoclinic S(0) phases were formed abiotically at low temperature in the presence of organics (Choudhury et al., 2013; Cosmidis et al., 2019; Guo et al., 2006; Moon et al., 2013). In microbial incubations, soluble organic compounds produced by *S. kujiense* were found to play an important role in the formation and stabilization of extracellular β-S_8_ particles in cultures and in spent media containing soluble organics (Cron et al., 2019). β-S_8_ was also previously described from a low-temperature natural environment and it was proposed that it was stabilized by its intimate association with organic matter (Lau et al., 2017).

Based on FTIR and XANES, the organic material associated with S(0) in the biofilms has a complex composition dominated by carboxylic acids, amides, aromatics, and aliphatic compounds (Figs. 6,8). It is important to note that the organic compounds we observed are not merely “associated”, but directly encapsulating S(0) particles (Fig. 7A,B). Consistent with this observation, previous work describing organics associated with S(0) formed in organomineralization experiments demonstrated the presence of carboxylic (Cosmidis et al., 2019) and amide groups (Cosmidis and Templeton, 2016) in organic envelopes closely encapsulating S(0) minerals. Pure cultures of the *Sulfurovum* relative *S. kujiense* also produced an amide-rich envelope around extracellular S(0) globules (Cron et al., 2019). Similarly, extracellular S(0) produced by *C. tepidum* was encapsulated in organic envelopes composed of proteins and polysaccharides (Marnocha et al., 2019).

Both organic envelopes around S(0) particles and the presence of the metastable allotrope β-S_8_ thus confirm an important role for organic-mineral interactions in extracellular S(0) mineralization. We therefore hypothesize that microbially derived extracellular organics (EPS) are critical for the formation and preservation of S(0) particles in the *Sulfurovum*-dominated biofilms at Frasassi, raising the interesting question below.

### 4.2 S(0) organomineralization and storage in biofilms: a new role for EPS?

Extracellular formation of biominerals in close interaction with organic polymeric structures has been documented for different types of systems, for instance precipitation of calcium carbonates in microbial mats (Dupraz et al., 2009) or on diatom EPS (Stanton et al., 2021), or iron-(oxyhydr)oxide mineralization on polymeric bacterial sheaths and stalks (Chan et al., 2009; Chan et al., 2011). While in some cases mineral precipitation on extracellular organics may be “unintended and uncontrolled” (Frankel and Bazylinski, 2003), in other cases specific organic structures are involved in directing extracellular biomineralization and this process plays a crucial role in metabolism, growth and/or survival (Chan et al., 2016; Miot et al., 2009).

In the Frasassi *Sulfurovum*-rich biofilms, EPS appear to influence S(0) formed as a result of microbial S-oxidation, through the stabilization of otherwise unstable mineral phases. Organics have been shown to favor the formation and stabilization of S(0) minerals through S(0) organomineralization (Cosmidis et al., 2019; Cosmidis and Templeton, 2016). S(0) organomineralization may occur with diverse types of organic molecules (e.g., amino-acids, sugars, humic acids; Cosmidis et al., 2019), as well as with soluble organic compounds produced by different types of bacteria (Cron et al., 2019). It remains to be determined whether specific organics are produced by the Frasassi biofilm microbial community to direct S(0) formation and stabilization.

Given that organic matter is energetically costly to produce (Amend et al., 2013; Jayathilake et al., 2017), particularly for autotrophs like *Sulfurovum* (LaRowe and Amend, 2015; LaRowe and Amend, 2016), we consider it unlikely that S(0) encapsulation and particle trapping in the biofilm is accidental. Holdfasts in colonizing *Sulfurovum* populations appear to be made of S(0) rather than organic matter, eliminating a potential role for EPS in biofilm attachment. These observations suggest that binding and encapsulating S(0) in EPS may represent a particular ecological strategy.

We can speculate about the ecological role that EPS and embedded S(0) particles might play in the survival and dispersal of *Sulfurovum* populations in the cave habitat. In this environment, *Sulfurovum*-rich biofilms form only in turbulent stream flows near the air-water interface, and are absent when appropriate support structures are unavailable in that position. The entrenched cave streams hosting the biofilms are subject to rapid, weather-related changes in water levels that frequently wash out or drown the biofilms. Thus cells that are producing S(0) under ideal conditions (Reaction 2) may experience rapid changes in O2 availability. Under O_2_ starvation conditions, bioavailable S(0) in the biofilm could enable survival and possibly even growth. Hamilton et al. (2015) noted that *Sulfurovum* metagenome-assembled genomes (MAGs) obtained from biofilms collected at the same sampling site ubiquitously contained genes that would allow H2 and formate oxidation with S(0) as an electron acceptor (Reaction 1).

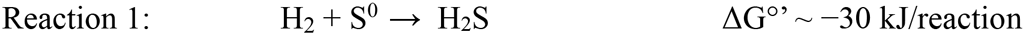

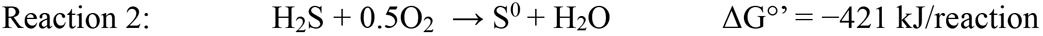

Although Reaction 1 yields much less energy than Reaction 2 under standard conditions (and likely also under *in situ* conditions), the ability to utilize stored S(0) may represent an important ecological advantage to the cave *Sulfurovum* populations, and capturing and storing S(0) in EPS may be an important element in the life strategy of this group. This scenario will be tested using metatranscriptomics in future work. If so, energy storage through stabilization of electron donors for energy metabolism may be a previously underappreciated role for EPS in biofilms.

EPS in the Frasassi biofilms described here appear to be responsible for at least two properties of the S(0) minerals: their crystal structure (with the stabilization of β-S_8_), and organic encapsulation. It is now well established that the properties of S(0) particles play a crucial role in their utilization by bacteria. For instance, particle size, surface area, and S^0^ composition and structure affect S(0) oxidation rates by *Thiobacillus albertis* (Laishley et al., 1986). *Allochromatium vinosum* grown on S(0) show a preference for polymeric sulfur over commercial crystalline S_8_, which they are unable to uptake (Franz et al., 2007). Preference for polymeric sulfur utilization over S_8_ was also evidenced in natural mats of chemotrophic S-oxidizers (Engel et al., 2007). Incubation experiments of natural freshwater communities with different sulfur sources showed a preference for the utilization of a reactive form of colloidal S(0) – possibly polythionates – over S_8_ (Findlay and Kamyshny, 2017). Recently, it was shown that *C. tepidum* can grow from the oxidation of its own biogenic S(0) globules but not from oxidation of commercial sulfur, crystalline S(0), or inorganically precipitated colloidal S(0), which is mineralogically very similar to biogenic S(0) globules but does not have an organic coating (Marnocha et al., 2016; Marnocha et al., 2019). It is possible that S(0) formation within EPS favors storage in a form more readily utilizable by the cells, by favoring metastable S(0) structures and by making S(0) particles hydrophilic, allowing interaction with the cell surface (Marnocha et al., 2019).

## 5 Conclusion

We studied a natural *Sulfurovum-*rich S(0)-producing biofilm to investigate processes that influence the precipitation and stabilization of extracellular S(0) in the environment. We found that sulfur particles in the biofilm are encapsulated within organic envelopes and that some of the particles have an unstable crystal structure (the high-temperature allotrope β-S_8_). These characteristics have also been observed in S(0) produced by organomineralization. Our results suggest that EPS within the biofilm stabilize S(0) particles, preventing their dispersal away from the biofilm and influencing their structural and surface properties. Future studies will be needed in order to determine the ecological importance of this process and its impact on biogeochemical sulfur cycling in the Frasassi cave and in other environments.

## 8 Author Contributions

B.C., J.L.M. and J.C. designed the study. B.C. and J.L.M. collected the samples, and B.C. and J.C. performed the analyses. All authors participated in data interpretation. B.C. wrote the manuscript with input from J.C. and J.L.M..

## 10 Acknowledgments

We thank C. Nims and M. Wetherington (Penn State University) for help with Raman analyses, and T. J. Zimudzi (Penn State University) for help with FTIR measurements. Use of the SSRL, SLAC National Accelerator Laboratory, is supported by the U.S. Department of Energy, Office of Science, Office of Basic Energy Sciences under Contract No. DE-AC02-76SF00515. We thank J. Wang for providing support on STXM beamline SM of the Canadian Light Source (CLS, Saskatoon, Canada). CLS is supported by the Canada Foundation for Innovation, Natural Sciences and Engineering Research Council of Canada, the University of Saskatchewan, the Government of Saskatchewan, Western Economic Diversification Canada, the National Research Council Canada, and the Canadian Institutes of Health Research. The authors are grateful to A. Montanari for providing facilities and laboratory space at the Osservatorio Geologico di Coldigioco in Apiro, Italy. Fieldwork was carried out in collaboration with C. Chan, C. Clark, and P. Henri with assistance from M. Mainiero, S. Mariani, and S. Recanatini.

