## Supplementary Material for "Organic stabilization of extracellular elemental sulfur in a *Sulfurovum*-rich biofilm: a new role for EPS?" (from another version of this preprint: version 1, 2021-06-03)

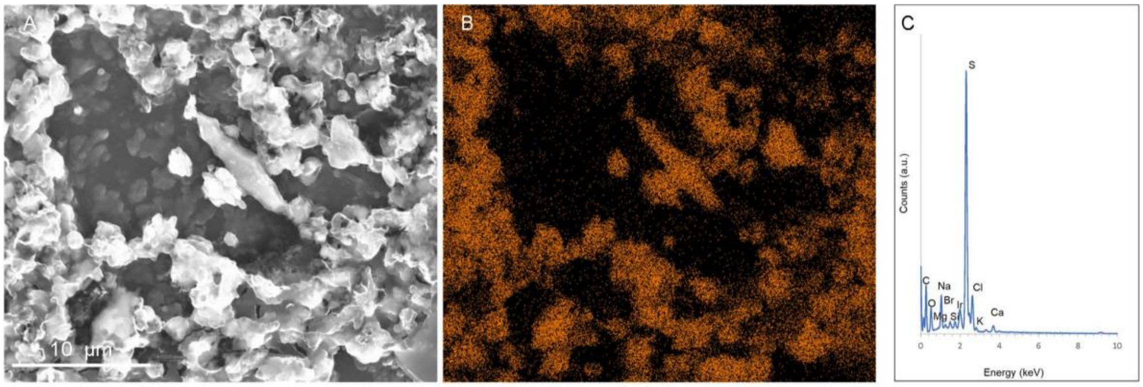


**Supplementary figure 1.** (A) SEM image of *Sulfurovum*-dominated streamer biofilm collected at Pozzo di Cristalli (PC1718). (B) EDXS map of the same area, showing sulfur distribution (orange). (C) EDXS spectrum. Additional elements indicate the presence of other minerals such as carbonates, quartz, and possibly clays.

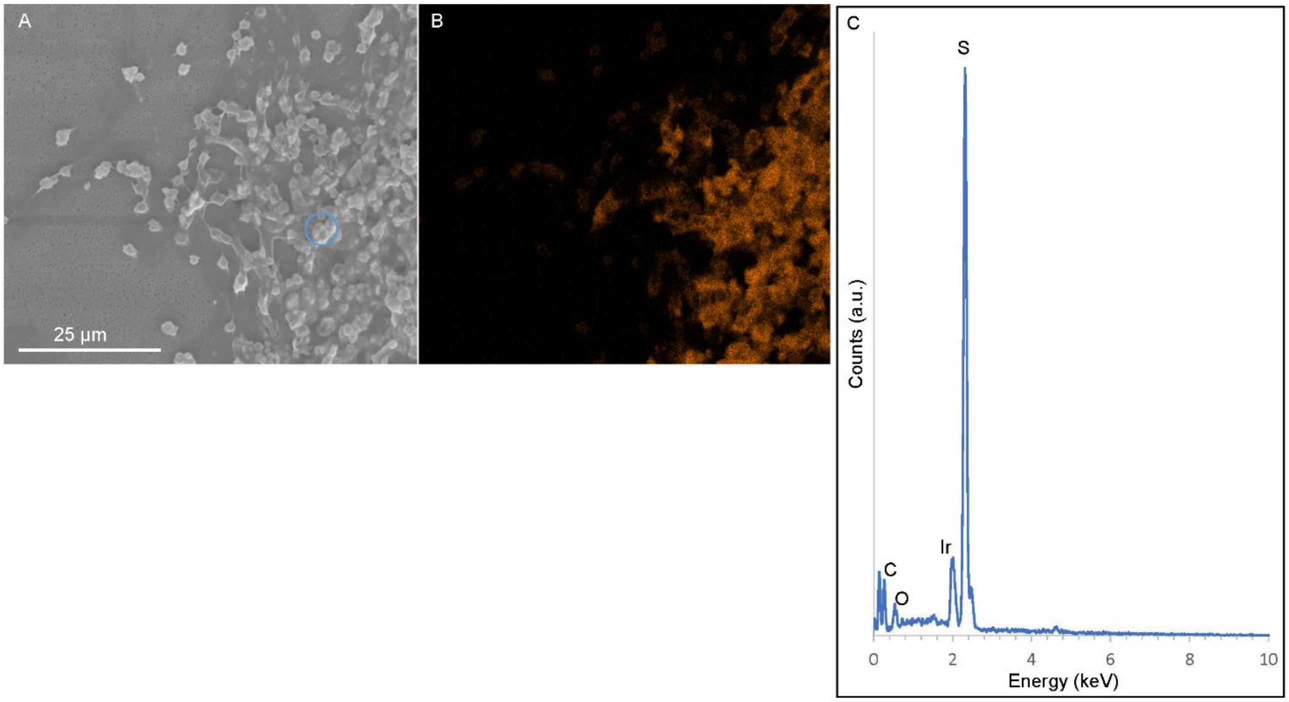


**Supplementary figure 2.** (A) SEM image of a *Sulfurovum*-dominated streamer biofilm collected at Pozzo di Cristalli (PC1718). The blue circle indicates the location where the EDXS spectrum in (C) was acquired. (B) EDXS map of the same area showing sulfur distribution (orange). (C) EDXS spectrum.

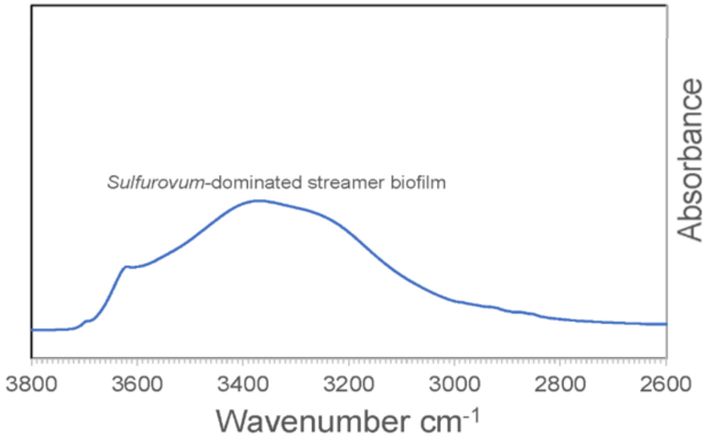


**Supplementary figure 3.** FTIR spectrum for a *Sulfurovum*-dominated streamer biofilm collected at Pozzo dei Cristalli (PC1718).

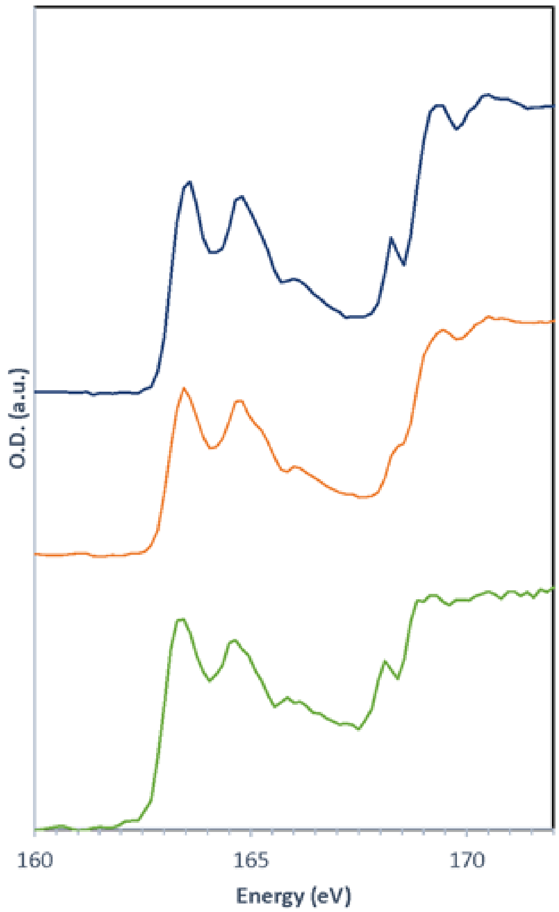


**Supplementary figure 4.** XANES spectra at the S L-edge obtained on particles from a *Sulfurovum*-dominated streamer biofilm (PC1718). All three spectra correspond to S(0).
